# GTEM cell design for controlled exposure to electromagnetic fields in biological system

**DOI:** 10.1101/2022.03.03.482821

**Authors:** J. Tao, Z.-G. Li, T.-H. Vuong, O. Kunduzova

## Abstract

This study describes a structure design of the Gigaherz Transverse Electromagnetic (GTEM) cell for electromagnetic tests in biological systems. Using electromagnetic simulation software we show that the optimization of final structure assures the criteria imposed on a low level of reflection towards the radio frequency generator, and a fairly uniform field in the test zone. A sensitivity study was also conducted when the cell was loaded with a few samples under test. We report that the level of reflection remains adequate as long as the test volume remains limited in relation to the size of the cell. The first estimation results on the SAR values are consistent with the distribution of the fields in the cell. A cell was fabricated with the design data, and tested for the reflection level. The measurements confirm the results of simulations.

## I. INTRODUCTION

With the rapid development of the social economy and modern science and technology, the application of radio frequency equipment is increasingly widespread, and the electromagnetic environment in which human beings is located is becoming more and more complex. According to recent studies, these electromagnetic waves will not only generate the EMI (Electromagnetic Interference) in some electronic devices, but also affect people’s health to a certain extent.

At present, the study of the electromagnetic environment covers two main aspects including EMC (Electromagnetic Compatibility) [1] with EMI and EMS (Electromagnetic Sensitivity Testing) issues and Bio-electromagnetic [2]. For the experiments in the above two cases, we all need a zero-interference environment to explore the relationship between the electromagnetic wave and the target devices/living body. When large object is to be studied under electromagnetic fields exposure on a laboratory scale, the GTEM cell can be optimal compromise between a Transverse Electromagnetic cell (TEM cell) and an anechoic chamber or a reverberation chamber.

In this article we present our work of designing a GTEM cell. After the estimation of the geometrical parameters of the GTEM cell and the introduction of these parameters into the electromagnetic simulation software HFSS, an optimization work was carried out in terms of the level of reflection towards the radiofrequency generator and the homogeneity of the electric fields in the working area. The influence of media under electromagnetic radiation will also be studied through the use of a number of biological samples. Results will be given on the level of reflection, on the distribution of the fields inside the cell and in the area of exposure. SAR estimates will also be given. Experiments on cardiac tissues are in progress, and will be presented later.

## II. CHOICE AND DESIGN OF EXPOSURE SYSTEM

### II.1. Background of GTEM Cell

GTEM cell is developed on the basis of the TEM cell. By generating uniform TEM electromagnetic waves inside the chamber, electromagnetic compatibility-related performance tests can be performed.

The early TEM was an open stripline, but it needed to be tested in the shielded room. Later, there was a closed-space TEM room, but the upper limit of the available frequency was limited to hundreds of megahertz due to the available space [3]. In order to solve the shortcomings of the lower upper limit frequency of TEM cells, in 1989, D. Hansen and D. Konigstein proposed GTEM [4].

### II.2. GTEM cell considerations

The cell to be manufactured must have a length not exceeding 1.4m, and must be powered by a female N-type coaxial connector, with a characteristic impedance of 50ohms.

The pyramidal structure will have a progressive taping starting from a symmetrical stripline of dimension slightly greater than that of the base plate of N connector, towards an asymmetrical stripline at the ending part, so that the large samples can be housed under the central conductor central. It is sought to maintain constant characteristic impedance throughout the structure, so that for a given transverse plane with respect to the pyramid, the equivalent capacitance is that of a 50ohms asymmetric stripline, with air between the central conductor and the metallic outer frame.

This capacitance will be given, according to [5], by the following relations

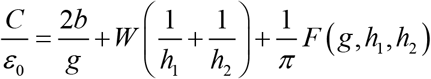

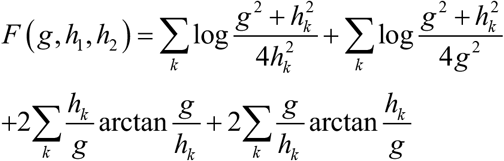

Here the central conductor has width *W*, and thickness *b. g* denotes the gap between central conductor and lateral walls, and *h*_1_ and *h*_2_ are distances between the central conductor and the upper and lower walls

A first set of GTEM cell parameter is deduced from these equations and the global size consideration.

### II.3. Optimization of GTEM Cell by HFSS

According to the working frequency of the signal source (a solid-state microwave source of 915MHz), the initial data has been introduced in a HFSS model. By tuning a certain number of parameter a GTEM cell with 1.4m length is obtained, as illustrated by Fig.1 and Fig.2.

**Fig. 1.**
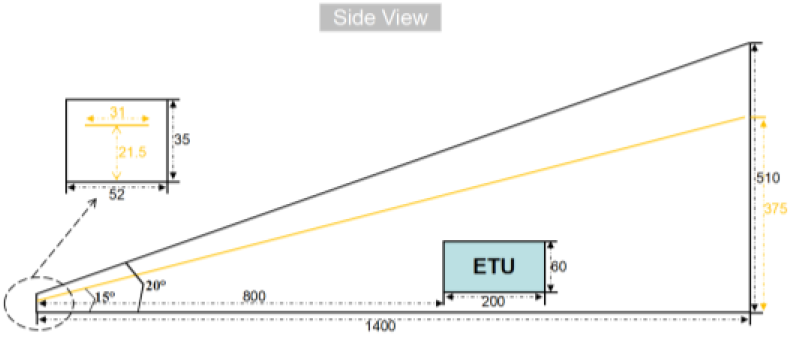
The side view of designed GTEM cell.

**Fig. 2.**
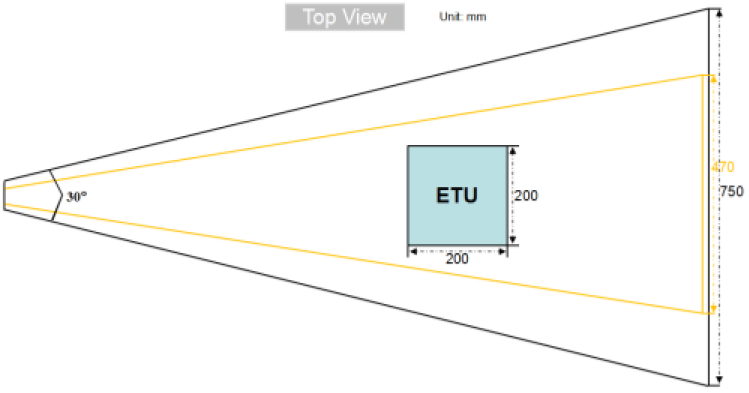
The top view of designed GTEM cell.

All sizes are summarized in Table I.

**Table 1.** Main parameters of GTEM cell(mm)

| Name | Len | FroW | FroH | EndW | EndH |
| --- | --- | --- | --- | --- | --- |
| Size | 1400 | 52 | 35 | 750 | 510 |
| Name | CopL | CopFW | CopFH | CopEW | CopEH |
| Size | 1395 | 31 | 21.5 | 470 | 375 |

In order to simplify the simulation, the N connector is not modeled in HFSS; in its place a symmetrical stripline of 50 ohms was used.

In the HFSS software, the provisions on the upper limit frequency that can be calculated is

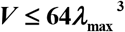

Where ***V*** is the volume of the model and *λ*_max_ is the upper limit frequency that can be calculated. According to this formula, the volume of this model is approximately 0.1785m3, so in the model simulation, the upper limit calculation frequency of this model is 6GHz, which meets the experimental requirements.

## III. NUMERICAL VERIFICATION OF DESIGNED GTEM CELL

Two criteria used in the design are now to be verified: the level of reflection towards the radiofrequency generator, and the uniformity of field in the area of exposure.

### III.1. Reflection of GTEM Cell

The results from optimized simulation model lead to the reflection coefficient of GTEM cell as shown in the Figure.3 below.

**Fig. 3.**
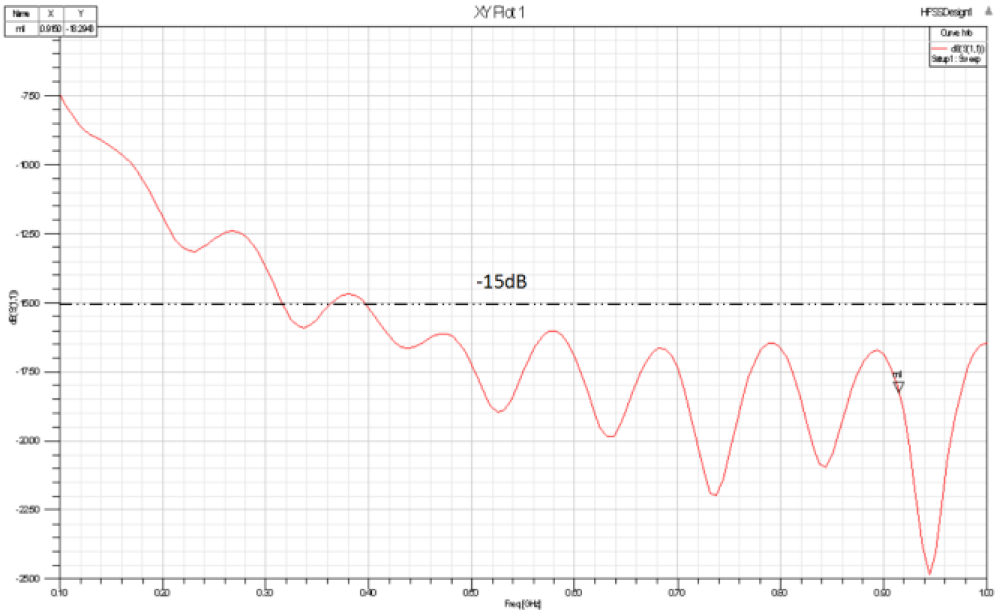
Reflection of designed GTEM cell.

**Fig. 4.**
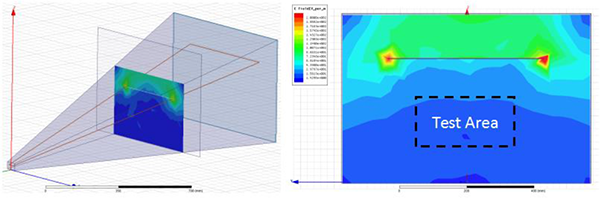
Field strength distribution at selected sections.

We can see that the reflection becomes lower than -15dB since 400MHz so that the low level is obtained for 915MHz, the working frequency.

### III.2. Field uniformity of GTEM Cell

GTEM cell as a test equipment requires that its internal field distribution must be very uniform [7]. The field strength distribution is shown in below.

As we can see, between the central conductor and the bottom wall, the internal spherical wave can be approximated as a plane wave, similar to the electromagnetic waves in the far field in free space, and the test area is completely contained within the uniform electromagnetic wave, responding to our requirement on electromagnetic wave exposure test.

### III.3. Influence of biological media

It is well known that living experiments must have an aqueous medium as a carrier to provide the necessary water, ions and nutrients.

In this first experiment, the biological environment will be represented by using three different concentrations of water as mediums, which are contained in the rectangular Equipment Under Test (EUT), as shown in Table.2. The EUT will be placed in the interior of the GTEM cell to study the relevant radiation characteristics, such as the reflection coefficient, Electric field strength, Specific absorption rate (SAR), etc.

**Table 2.** Conductivity of three different water.

| Mediums | Distilled water | Fresh water | Sea water |
| --- | --- | --- | --- |
| Conductivity | 0.0002 | 0.01 | 4 |

The results from optimized simulation model lead to the reflection coefficient of GTEM cell as shown in the Fig.5 below.

**Fig. 5.**
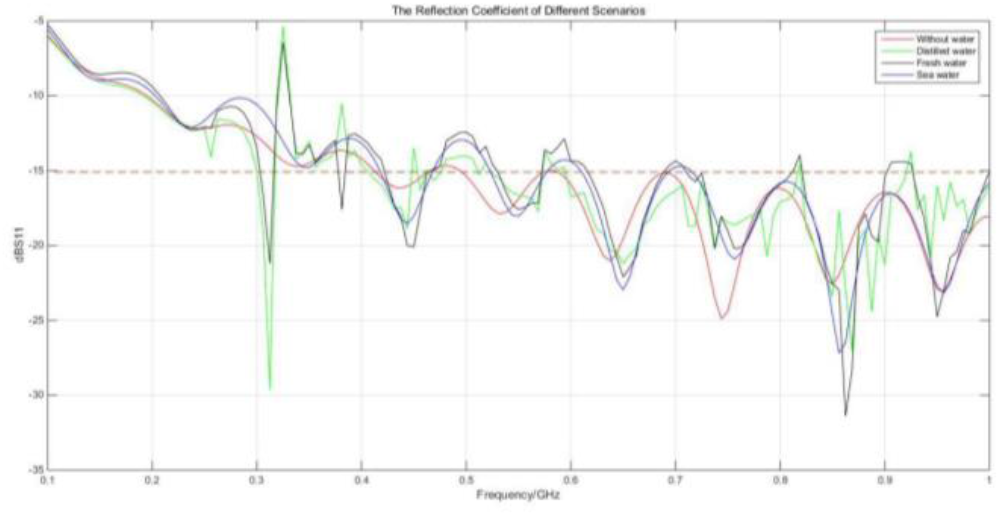
Reflection of GTEM cell with EUT.

Since the EUT is relatively small in size (200 mm x 200 mm x 60 mm), the effect on the reflection coefficient is very low. Therefore, the above four curves have a substantially identical shape, and between 300 MHz and 1 GHz, the reflection level is also substantially below -15 dB.

## IV. NUMERICAL RESULTS ON SAR ESTIMATION IN EUT

In general, various tissues of an organism are depleted media, so electromagnetic field acting on a biological medium generates an induced current, resulting in absorption and dissipation of energy.

The SAR is used to characterize this process, it is defined as the total electromagnetic energy that a biological tissue or organism can absorb per unit mass when exposed to an electromagnetic field, in units of W/Kg.

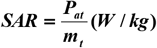

Where *P*_*at*_ is the total power absorbed by the organism and *m*_*t*_ is the total mass of the organism.

In actual calculations, the radiated power is not easily obtained directly, so the electric field strength ***E*** can be used to represent the SAR. The relationship between the SAR at a certain point and the field strength at the point is

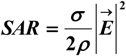

Where ***σ*** is the conductivity of the biological tissue at the location taken and ***E*** is the total electric field strength at that point [9].

In the HFSS simulation experiment, by setting the local SAR, we can directly observe the SAR distribution. As a comparison, in this experiment we set up a EUT with only vacuum medium in HFSS, and theoretically its SAR value should be zero. Therefore, the simulated SAR distribution in those four different mediums scenarios is shown in the Fig.6.

**Fig. 6.**
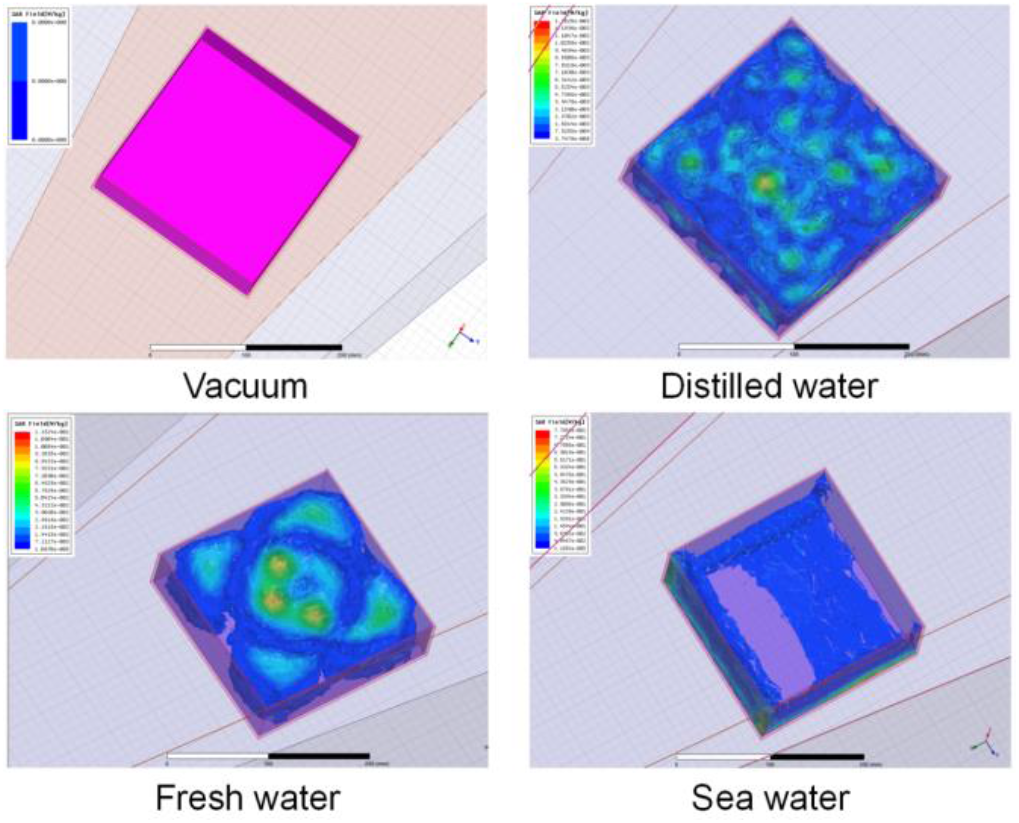
SAR distribution of different scenarios.

The distribution of SAR is positively correlated with the distribution of electric field strength not given in this paper. The maximum value of SAR is shown in Table 3 below.

**Table 3.** Maximum SAR for different scenario (W/kg)

| Mediums | Distilled water | Fresh water | Sea water |
| --- | --- | --- | --- |
| Maximum | 0 | $1.26 \times 10^{-2}$ | $1.15 \times 10^{-1}$ |

From this table it is easy to conclude that the higher the conductivity of the medium, the higher the maximum SAR, which means that the electromagnetic energy is more concentrated in one part of the medium. As shown in seawater medium, electromagnetic waves have little effect on the inside and the back and the propagation attenuation is very intense.

## V. CONCLUSION

This study introduces a structure design of GTEM cell for electromagnetic test in biological systems. Using electromagnetic simulation software we demonstrate that the optimization of final structure satisfies the criteria calibrated on a low level of reflection towards the radio frequency generator, and a fairly uniform field in the test area. A sensitivity study was also conducted when the cell was loaded with the samples under test. We demonstrate that the level of reflection remains adequate as long as the test volume remains limited in relation to the size of the cell. The first estimation results on the SAR values are consistent with the distribution of the fields in the cell.

A cell was fabricated with the design data, and tested for its reflection level. The measurements confirm the results of simulations. Tests on a series of biological samples are in progress.

